# Self-targeting spacers reveal new functions of CRISPR systems

**DOI:** 10.1101/2023.07.26.550751

**Authors:** Miguel Angel Tangarife Cardona, Juan Camilo Arboleda Rivera

**Affiliations:** Grupo de Bacteriología Agrícola y Ambiental, Instituto de Biología, Universidad de Antioquia UdeA, Medellín, Colombia; Grupo de Fundamentos y Enseñanza de la Física y los Sistemas Dinámicos, Instituto de Biología, Facultad de Ciencias Exactas y Naturales, Universidad de Antioquia UdeA, Medellín, Colombia

**Keywords:** CRISPR, Self-targeting spacers, Autoimmunity, Microbial genetics, Computational biology

## Abstract

The CRISPR systems enable bacteria and archaea to defend from bacteriophages or mobile genetic elements by inserting portions of the DNA of these elements into its own genome in sequences known as spacers that will later trigger the complementarity-based degradation of invading sequences. The presence of self-targeting spacers is widespread in prokaryotes; however, its functional role is still unclear. In this study, we analyzed self-targeting spacers of CRISPR systems and found a high presence of membrane proteins, aminoacyl-tRNA synthetases and ATP-binding proteins. This is a novel report that supports other research linking CRISPR systems to membrane proteins and could explain the reported relationships between antibiotic resistance and presence of CRISPR systems.

## Introduction

Clustered Regularly Interspaced Short Palindromic Repeats (CRISPR) and CRISPR associated proteins (Cas) are a molecular system that provides adaptive immunity in bacteria and archaea. In a process known as adaptation, small pieces of DNA (referred to as protospacers) such as that coming from phages are processed by Cas proteins and inserted in the prokaryotic genome within the CRISPR array. Those inserted pieces are separated by repetitive sequences and are known as spacers, which are transcribed and processed to target complementary sequences and induce its degradation by Cas proteins ^1^. While the majority of spacers are complementary to phage genomes or mobile genetic elements such as plasmids, self-targeting spacers have been reported since the beginning of the characterization of CRISPR sequences and have raised the question of whether they are mistakes of the CRISPR adaptation module leading to autoimmunity or if they have any function ^2^.

Although the functional relevance of self-targeting spacers has remained elusive, there is increasing evidence that self-targeting spacers are not mere accidents, but can harbor alternative functions ^3^, including regulation of gene expression ^4,5^.

In this study, we analyzed a database of self-targeting spacers and performed a functional characterization of its protospacers and found a link between the CRISPR system and specific groups of cellular proteins such as membrane proteins.

## Results

The original database consisted in genomic coordinates for self-targeting spacers derived mainly from ^6^ and some others from ^2^ and ^7^. This database contained information from 1634 species belonging to 278 different prokaryotic families (Figure 1), 259 of which were bacteria and 19 archaeal families.

**Figure 1.**
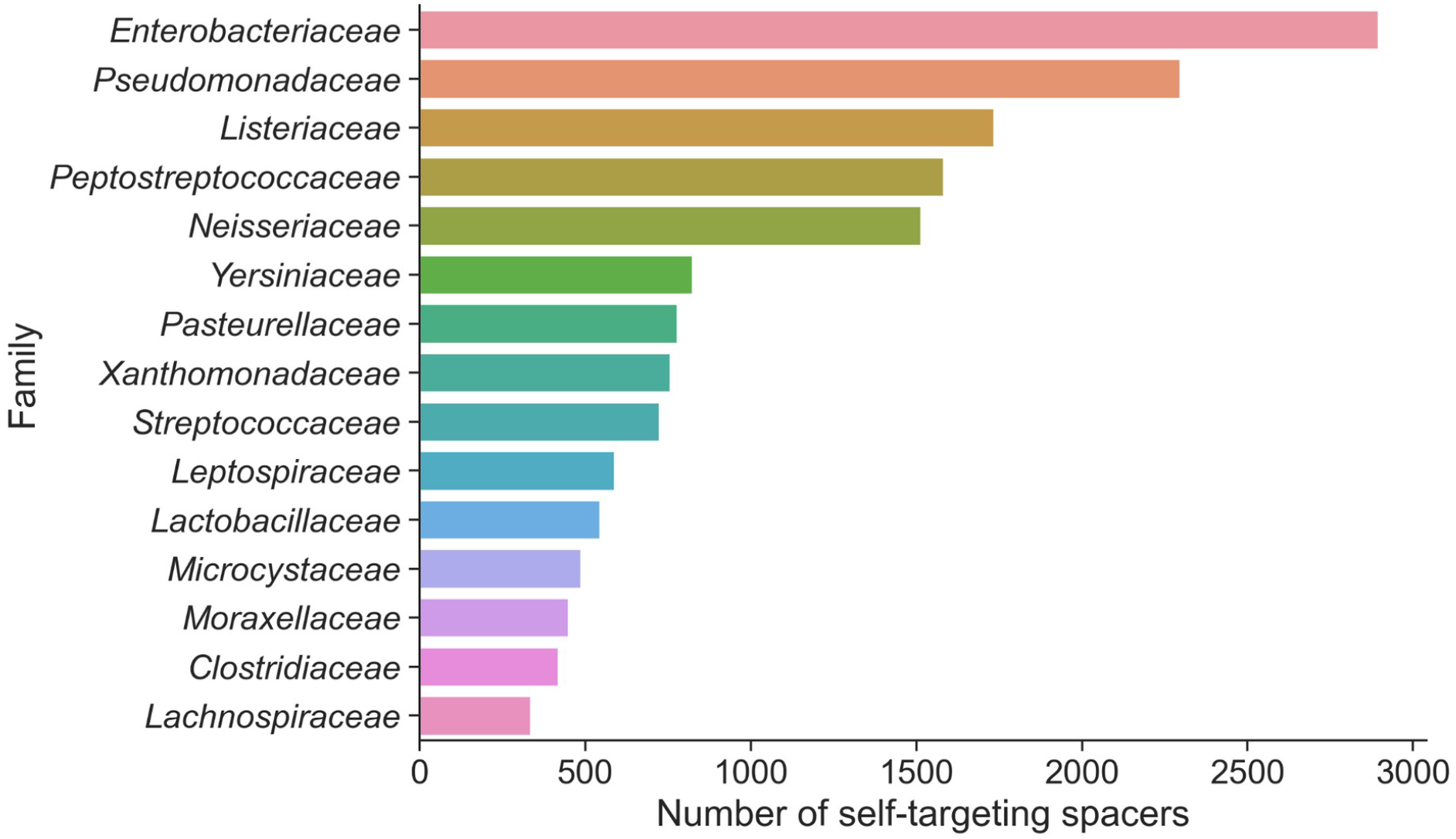
Representation of different families in the self-targeting spacers database.

Out of 22110 self-targeting spacers, 12428 were found to target known gene products and 1940 were found to target hypothetical proteins. 3592 spacers did not target known genes and no information could be retrieved from the NCBI for the remaining spacers.

In a previous analysis, Shmakov et al. ^8^ studied genes that were functionally linked to CRISPR systems by neighborhood analysis and found genes encoding proteins such as CARF domain-containing proteins, Lon family proteases and the CorA magnesium channel. We sought if those proteins were also targeted by self-targeting spacers in order to further confirm this association. We didn’t find CARF proteins but we found Lon protease targeting spacers in *Yersinia kristensenii, Enterobacter cloacae* and *Gilliamella apicola*, and CorA protein was found as a target in *Chloroflexi* sp. and nine different *Listeria monocytogenes* strains. Similarly, Ou et al. ^9^ observed a DedA protein coding sequence near the CRISPR locus in *Bifidobacterium*, and we found this protein as target of spacers in *Acetonema longum, Streptomyces* sp. and *Pseudomonas aeruginosa*.

ABC-transporters are the self-targeting spacers found in most species (Table 1), and were found in 74 different families, with a total of 165 different species having spacers targeting ABC domain containing proteins. Aminoacyl tRNA synthetases were another common group of protospacers, being alanine-tRNA ligase the most common of this group (Figure 3 and Table S1). Phage associated-proteins such as phage tail tape measure protein were also common, and come probably from prophages ^10^.

**Table 1.**
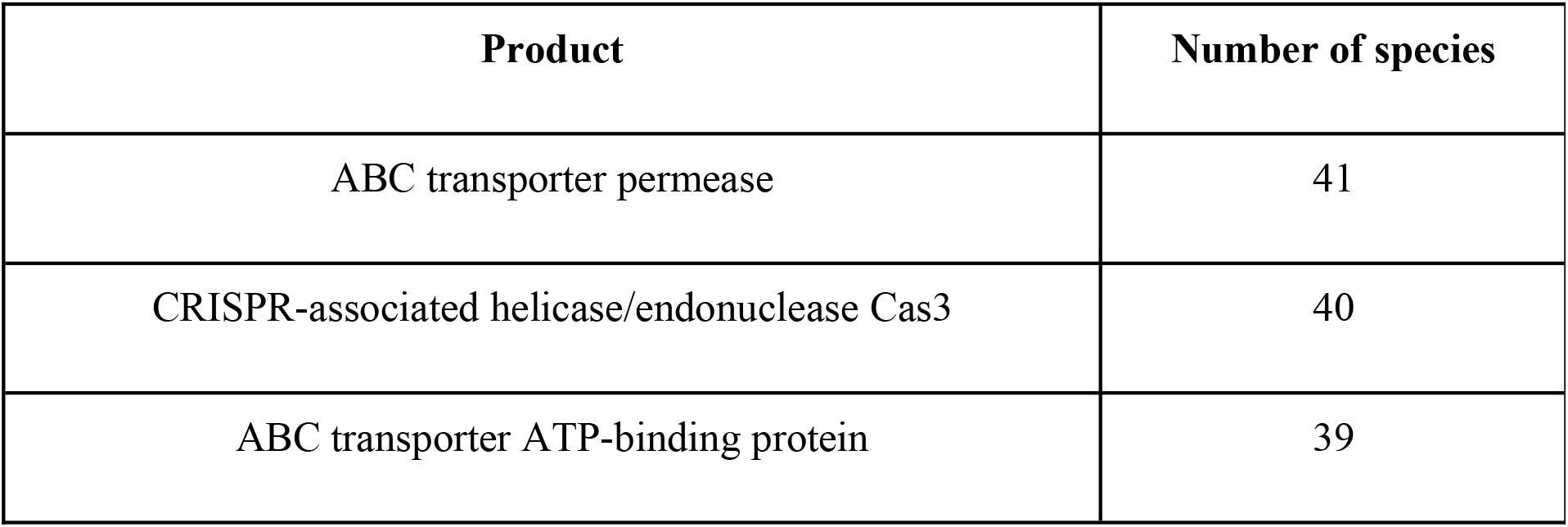
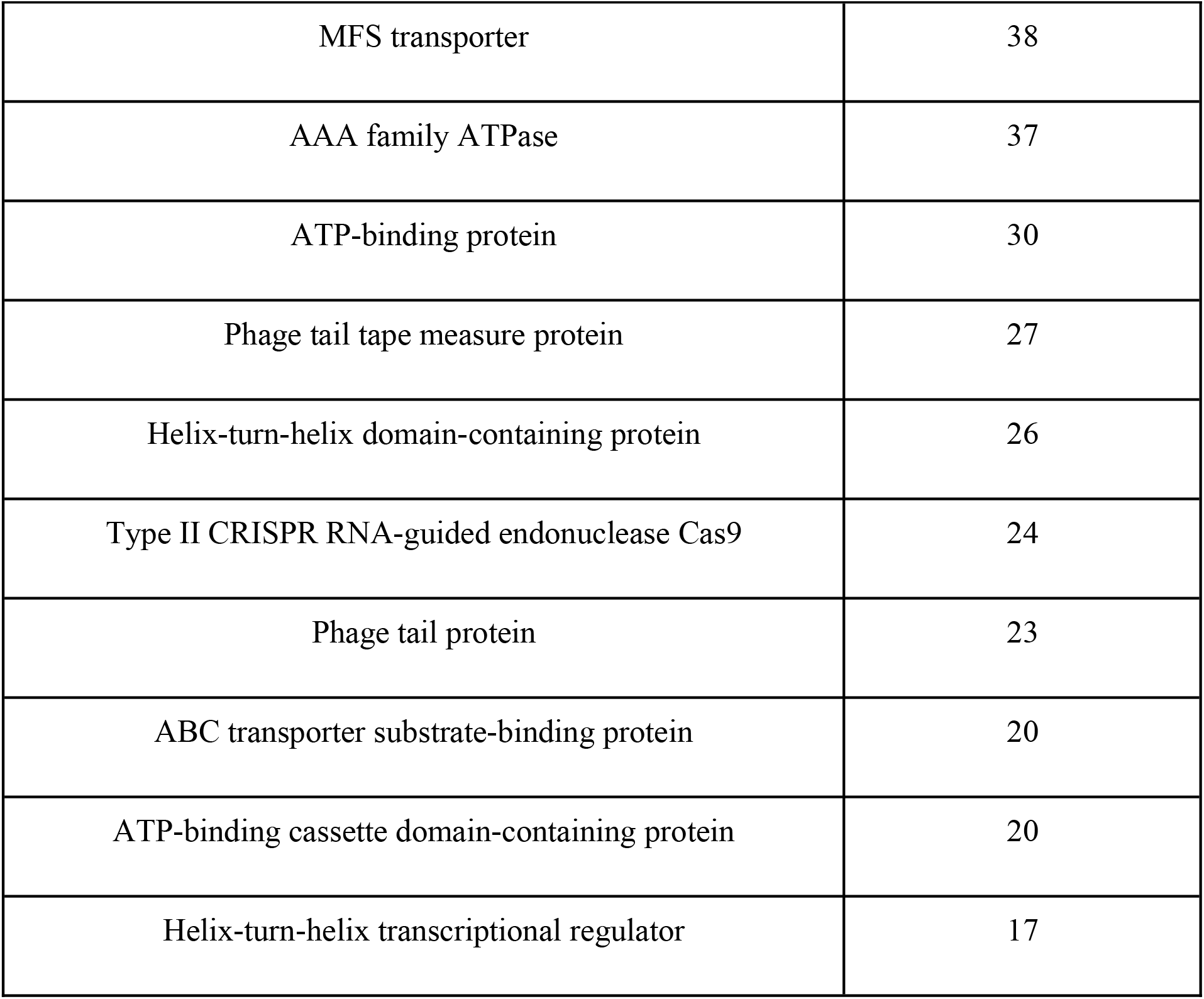
Most abundant gene products targeted by spacers in different prokaryotic species. Note the high presence of ATP-binding proteins.

| Product | Number of species |
| --- | --- |
| ABC transporter permease | 41 |
| CRISPR-associated helicase/endonuclease Cas3 | 40 |
| ABC transporter ATP-binding protein | 39 |
| MFS transporter | 38 |
| AAA family ATPase | 37 |
| ATP-binding protein | 30 |
| Phage tail tape measure protein | 27 |
| Helix-turn-helix domain-containing protein | 26 |
| Type II CRISPR RNA-guided endonuclease Cas9 | 24 |
| Phage tail protein | 23 |
| ABC transporter substrate-binding protein | 20 |
| ATP-binding cassette domain-containing protein | 20 |
| Helix-turn-helix transcriptional regulator | 17 |

We analyzed GO Process terms associated with targets of self-targeting spacers (Fig. 2). Maintenance of CRISPR repeat elements (GO:0043571) was the most common term, indicating a possible self-regulatory mechanism mediated by self-targeting spacers in CRISPR systems. Viral process (GO:0016032) was associated with 179 spacers and was expected to be a common term due to the canonical function of CRISPR systems.

**Figure 2.**
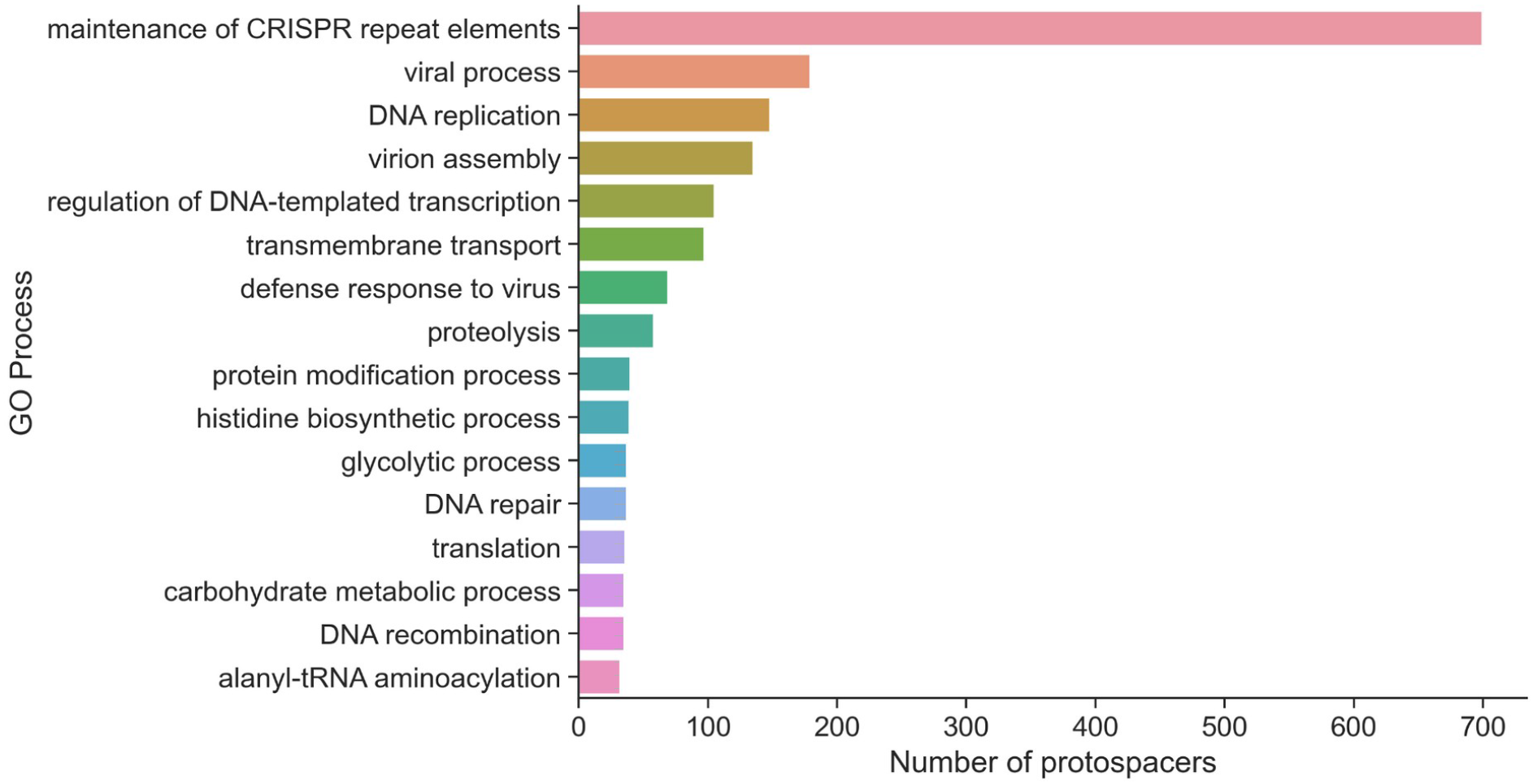
GO Processes associated with protospacers. Maintenance of CRISPR repeat elements is the most common GO process indicating a self-regulatory function of CRISPR spacers. It is interesting to note the high representation of the transmembrane transport process.

**Figure 3.**
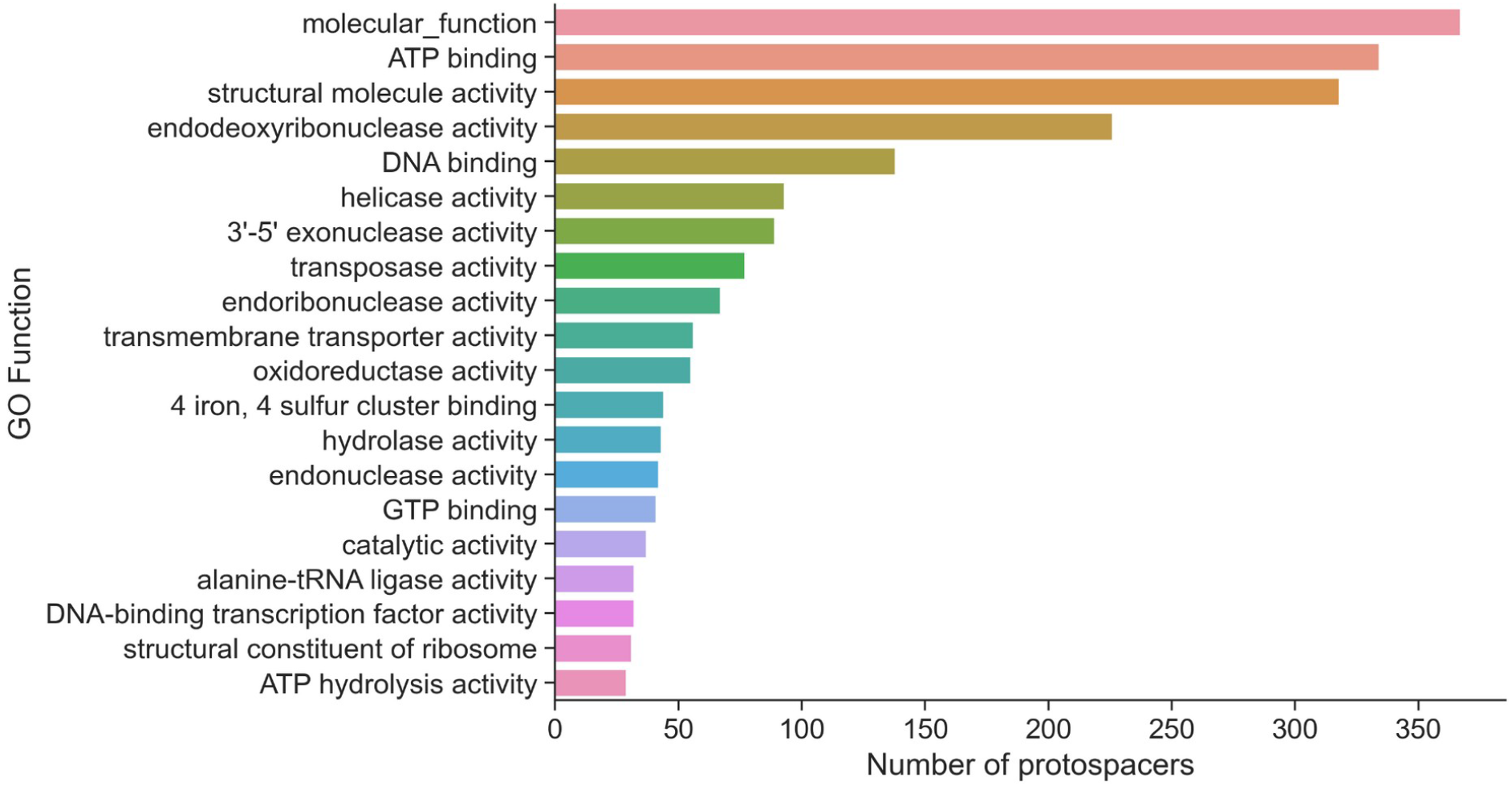
GO Functions associated with protospacers.

**Figure 3.**
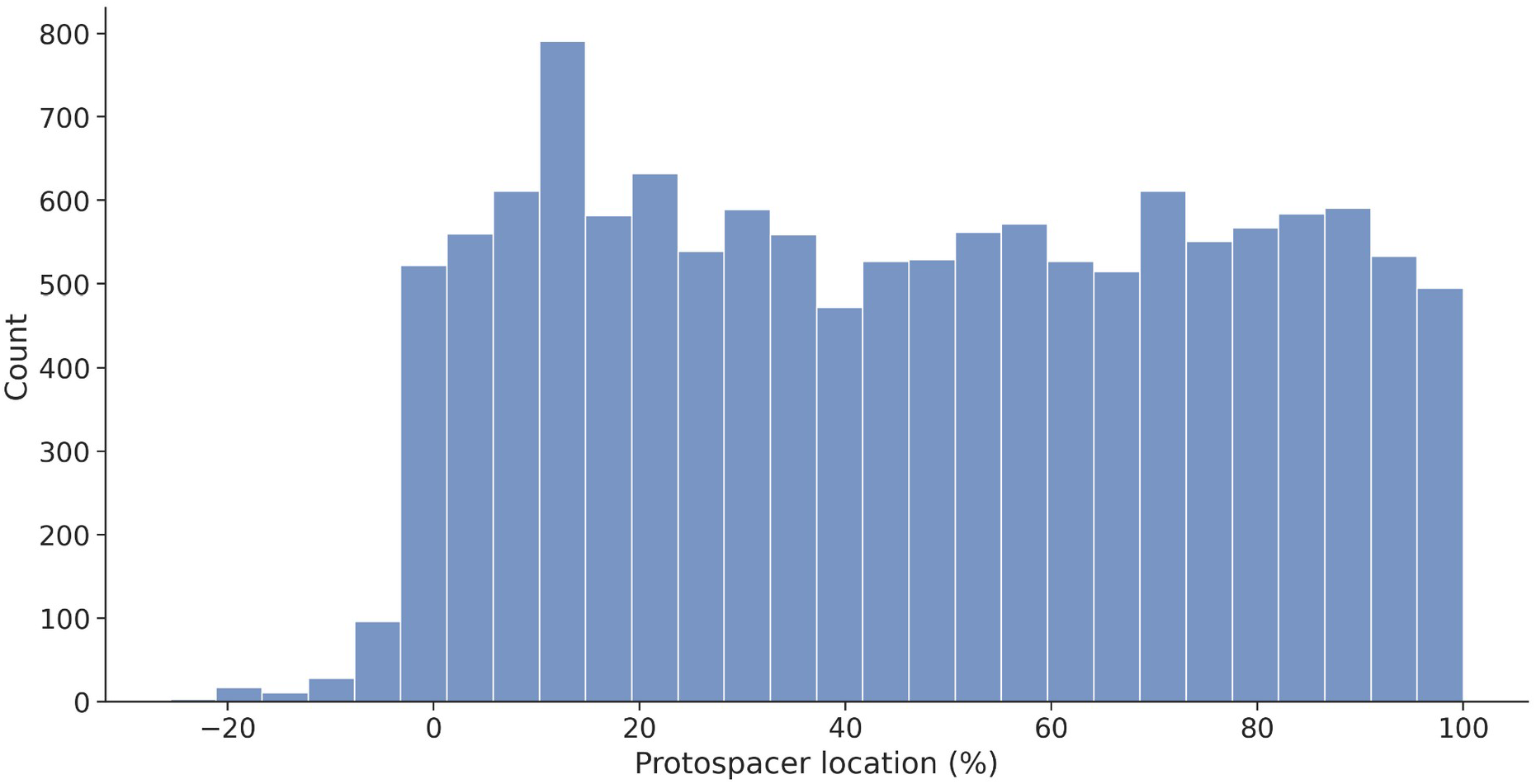
Protospacer location as percentage of the corresponding coding sequence. Protospacers are almost uniformly distributed along the coding sequence, although they are more common towards the 5’-end. Protospacers located before the translation start site are uncommon.

**Figure 4.**
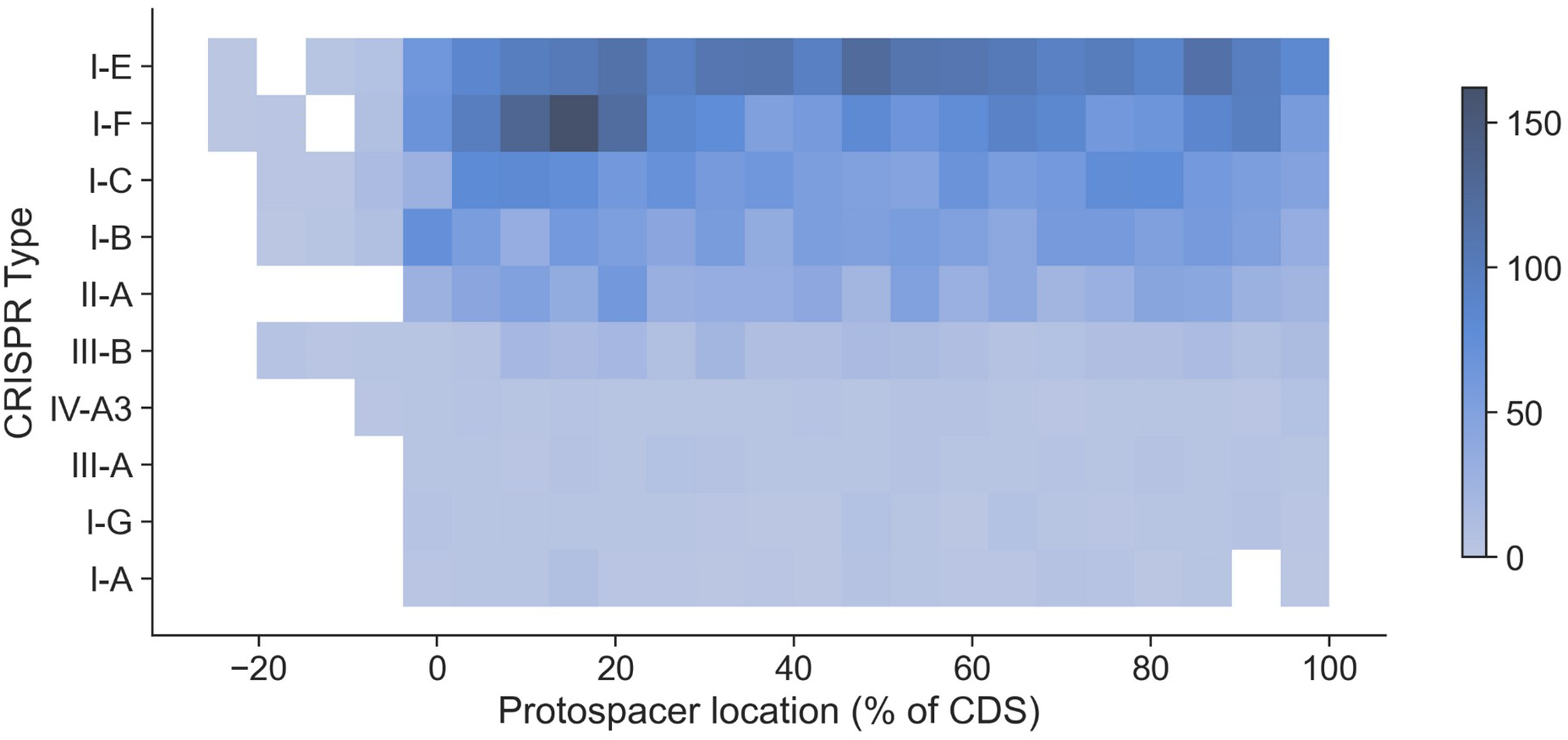
Protospacer location as percentage of coding sequence in the most abundant CRISPR Types. Most protospacers are uniformly distributed along the coding sequence; however, protospacers associated with CRISPR type I-F seem to be more common at the 5’-end.

We sought to classify each spacer according to its CRISPR type. We found that most spacers belonged to CRISPR systems of type I-E, I-F and I-C (Table 2); however, this can be explained by the abundance of Enterobacteria in the database, in which those CRISPR systems are most common.

**Table 2.** Main CRISPR types associated with self-targeting spacers. CRISPR I-E and I-F were the most commonly associated with self-targeting spacers, although its high abundance can be explained by the enrichment of *Enterobacteriaceae* in the database. Note the apparently self-regulatory function of each CRISPR system.

| CRISPR Type | Count | Main targeted product |
| --- | --- | --- |
| I-E | 2818 | CRISPR-associated helicase/endonuclease Cas3 |
| I-F | 2563 | Type I-F CRISPR-associated protein Csy1 |
| I-C | 2120 | Type I-C CRISPR-associated protein Cas8c/Csd1 |
| I-B | 1394 | CRISPR-associated helicase/endonuclease Cas3 |
| II-A | 1014 | Type II CRISPR RNA-guided endonuclease Cas9 |
| III-B | 537 | Type III-B CRISPR-associated protein Cas10/Cmr2 |
| IV-A3 | 172 | ATP-dependent helicase |
| III-A | 130 | (S)-acetoin forming diacetyl reductase |
| I-G | 104 | Type I-U CRISPR-associated RAMP protein<br>Csb1/Cas7u |
| I-A | 98 | ABC transporter permease |

As many protospacers were found to be annotated as hypothetical proteins, we performed a characterization *in silico* in order to infer its function. Functional prediction showed that 580 proteins matched the membrane term (GO:0016020) in the Cellular Component domain, while only 8 matches were found against metabolic process term (GO:0008152) in the Biological Process domain and 8 against catalytic activity (GO:0003824) in the Molecular Function domain.

The scan against the PROSITE database yielded 91 hits in 71 sequences. Of these hits 44 corresponded to a prokaryotic membrane lipoprotein lipid attachment site profile (PROSITE accession PS51257), indicating again a close association between self-targeting spacers and membrane proteins. The second most abundant motif had only 5 hits and was a TPR repeat region circular profile (accession PS50293). Analysis with Interproscan corroborated these results, showing hits against twin-arginine translocation (Tat) signal profile (S1 File).

## Methods

### Self-targeting spacers database

For this study, we used the CRISPRminer Self-Targeting database ^11^, which is based on the studies of Rauch et al., (2017); Stern et al., (2010) and Watters et al., (2018).

The CRISPRminer Self-Targeting database was downloaded on 2022-10-30. Additional information about self-targeting spacers was obtained from NCBI databases using Biopython’s Entrez module ^12^. Data analysis was performed using pandas ^13^.

### CRISPR typing

CRISPR type was predicted by inspecting the genomic region between 9000 bp upstream of the spacer start and 9000 bp downstream the spacer end using CCTyper version 1.7.1 ^14^.

### Function prediction of hypothetical proteins

We downloaded amino acid sequences from 1469 self-targeted hypothetical proteins from the NCBI’s Protein database. Functional prediction of hypothetical proteins was performed with DeepGOPlus v1.0.1 ^15^ with a threshold value of 0.3. Analysis of GO terms was performed using GOATOOLS ^16^. We also performed a scan of these sequences using Interproscan ^17,18^ and the ScanProsite tool excluding motifs with a high probability of occurrence ^19^.

### Protospacer location

In order to determine the location of protospacer inside coding sequences, we retrieved the coding sequence start (CDS_start_) and the coding sequence end (CDS_end_) from NCBI Nucleotide database. Then, we calculated the protospacer location (ps_loc_) by using the protospacer start (ps_start_) with the following equation:

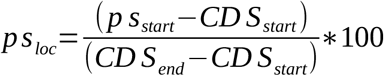

## Data and Material availability

Source code of the analysis and data used in this study are available from https://github.com/Nesper94/CRISPR-self-targeting.

## Discussion

It is remarkable to note that many CRISPR associated proteins were also found as protospacers (e.g., Lon proteases), indicating a functional link that remains to be discovered or a possible autoregulatory mechanism.

Self-targeting spacers leading to DNAse activity are likely to be harmful for the cell; however, if those spacers trigger RNAse activity or translation inhibition, the effect would be the regulation of gene expression. Auxiliary RNA-binding proteins may exist that fulfill this role, or known Cas proteins may have RNA-binding domains as is the case for Cas9 ^5,20^.

Type I CRISPR systems have been reported to induce cleavage of DNA targets; however, the presence of self targeting spacers directed to genes that are functionally related to infection processes in these systems indicates that they could have auxiliary functions regulating gene expression, maybe by binding to mRNAs and inhibiting translation or by binding directly to the gene and avoiding transcription. Moreover, it has been recently reported that the Type IV-A CRISPR-Cas system is capable of interference without involving protospacer degradation ^4^; however, these authors reported cleavage of protospacer for the Type I-C CRISPR-Cas system, while this was one of the most abundant CRISPR systems in the self-targeting spacers database. It is possible the existence of a mechanism that allows distinguishing between self and non-self protospacers in order to prevent self DNA degradation.

### Aminoacyl-tRNA synthetases

Another interesting aspect is the recurrent association between CRISPR and aminoacyl-tRNA ligases. For example, Aklujkar & Lovley ^21^ reported that CRISPR interfered with histidyl-tRNA ligase in *Pelobacter carbinolicus* and Ou et al. ^9^ found an isoleucine-tRNA ligase gene near the CRISPR locus of several strains of *Bifidobacterium*.

In our study, alanine tRNA ligase and leucine tRNA ligase were the most targeted aminoacyl tRNA synthetases, which could be explained by the fact that these are common amino acids in proteins, and thus targeting these enzymes could be an efficient way to shutdown protein synthesis when bacteria are infected with a virus and the CRISPR system is active.

### Membrane proteins

Previous analysis of CRISPR systems and CRISPR associated proteins has revealed a mysterious connection between CRISPR and membrane proteins ^8,22–24^, and our results support this association.

The regulation of membrane proteins by CRISPR could assist different functions; for example, it could be ligated to immune evasion in pathogenic bacteria as has been shown in *Campylobacter jejuni* and *Francisella novicida* ^5,25^. Moreover, membrane proteins can also be necessary for phage infection ^26^, so one possibility is that some self-targeting spacers are directed against those proteins, as is the case of TolC. In this way, the CRISPR systems could degrade invading nucleic acids and at the same time prevent infection by reducing the expression of receptors needed for phage entry. For example, the membrane-anchored protease FtsH was shown to be implicated in phage attachment ^27^, and was also found as target of self-targeting spacers in *Klebsiella pneumoniae, Nitrosococcus halophilus, Campylobacter coli, Neisseria sicca* and *Xenorhabdus ehlersii*. However, there seems to be another yet unknown functional aspect linking CRISPR and the cell membrane, as other types of membrane proteins such as those related to the Tat pathway have found to be related with the CRISPR system ^23^.

In a previous study, Shah et al. ^22^ identified genes associated with type III CRISPR systems and found a cluster of genes encoding ABC-transporters but disregarded this result as a spurious association. However, like this study, we also found AAA ATPases and ATP-binding proteins well represented in self-targeting spacers, validating the functional association of these components with CRISPR systems (Table 1). This result, together with the others we have presented here, reinforces the idea that self-targeting spacers are indeed in some way functionally linked ^3^ and are not mere errors leading to autoimmunity as had been previously believed ^2^.

Moreover, ABC transporter genes have been found physically linked to CRISPR-Cas locus ^9^. It is very interesting that relevant protospacers identified in this study were also found in the study of Ou et al. ^9^, it is the case of LysR family transcriptional regulator, helix-turn-helix domain-containing protein and MFS transporters.

Additionally, ABC transporters were found to be upregulated in a CRISPR-Cas knockout mutant ^28^. This regulation of ABC transporters by CRISPR systems could explain some observations related to antibiotic resistance in bacteria. In particular, some researchers have reported that resistance in bacteria is negatively correlated with the presence of CRISPR systems ^29–33^. This can be explained in part given that CRISPR gives immunity against mobile genetic elements such as plasmids that can carry resistance genes; however, at the light of our findings, we propose that bacterial resistance could be also linked to the presence of spacers targeting ABC transporters in CRISPR systems, although the functional or evolutionary relationship between these systems are not yet known to our knowledge.

Quorum sensing related proteins such as LuxR, LasR and AbaI seem to be another common target of self-targeting spacers ^3,28,34^. It is possible that disrupting biofilm formation to avoid phage propagation is the selection pressure causing this phenomena. Nevertheless, the functional link between CRISPR-Cas, ABC transporters, quorum sensing and other membrane components remain to be elucidated and further experimental confirmation is required.

In summary, this study highlights the importance of studying self-targeting spacers and supports the recent associations found between CRISPR systems and membrane components, an aspect of CRISPR biology that requires closer attention. It also highlights the fact that, while non canonical functions of CRISPR are not something new, they may be more common in prokaryotes than previously believed.

## Acknowledgments

We would like to thank Arley Calle Tobón for helpful discussions and critical reading of this manuscript. The computational resources used in this work were provided by the FEnFiSDi group computational cluster.

## Author Contributions

M.A.T.C. and J.C.A.R. contributed equally to this work.

## Additional Information

Supplementary Information is available for this paper.

## Supplementary material

**Table S1.**
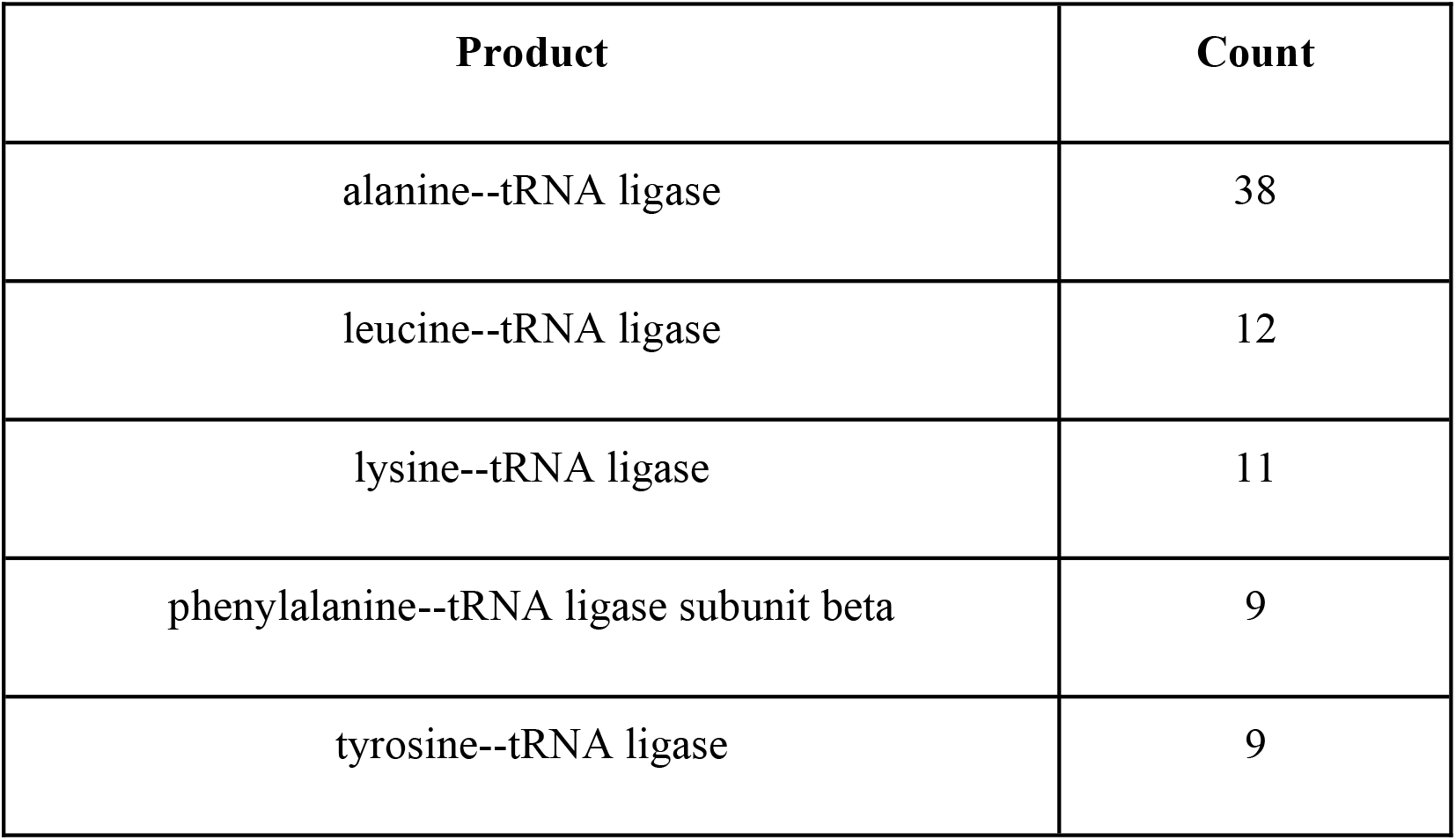
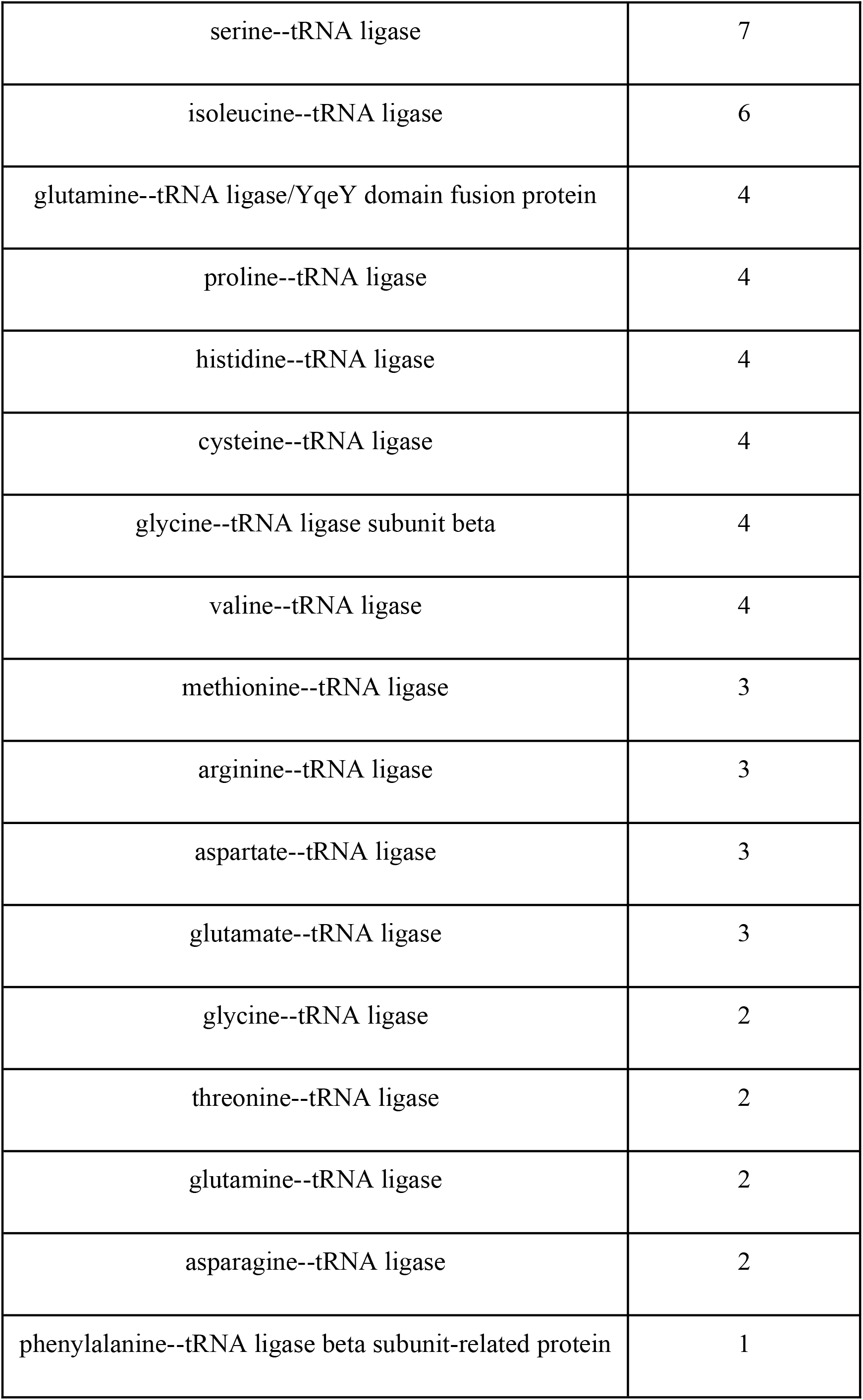
Abundance of self-targeting spacers directed to aminoacyl-tRNA ligases.

